# Global genomic analysis reveals the genetic origin and secondary invasion of fall armyworm in the Eastern hemisphere

**DOI:** 10.1101/2022.09.16.508260

**Authors:** Lei Zhang, Zaiyuan Li, Yan Peng, Xinyue Liang, Kenneth Wilson, Gilson Chipabika, Patrick Karangwa, Bellancile Uzayisenga, Benjamin A. Mensah, Donald L. Kachigamba, Yutao Xiao

## Abstract

The major plant pest fall armyworm (FAW), *Spodoptera frugiperda*, is native to the Americas and has colonized African and Asian countries in the Eastern hemisphere since 2016, causing severe damage to multiple agricultural crop species. However, the genetic origin of these invasive populations require more in-depth exploration. We analyzed genetic variation across FAW genomes of 153 newly sequenced individuals from Eastern hemisphere and 127 individuals mostly originating from the Americas. The global genetic structure of FAW shows that the FAW in American has experienced deep differentiation, largely consistent with the Z-chromosomal *Tpi* haplotypes commonly used to differentiate “corn-strain” and “rice-strain” populations. Results indicate that the invasive Eastern hemisphere populations are different from the American ones and have relatively homogeneous population structure, consistent with the common origin and recent spreading from Africa to Asia. Our analyses suggest that north-and central American “corn-strain” FAW are the most likely sources of the invasion into the Eastern hemisphere. Furthermore, evidence based on genomic, transcriptomic and mitochondrial haplotype network analysis suggest that there has been an earlier independent introduction of FAW into Africa that introgressed into the recent invasive population.

## Introduction

The fall armyworm (FAW), *Spodoptera frugiperda* (J.E. Smith), is a migratory pest native to tropical and subtropical regions in the Western hemisphere (Sparks 1979; Johnson 1987; Mitchell et al. 1991). Since the first detection of outbreaks in Africa in 2016, it has experienced an incredibly rapid expansion across the Eastern hemisphere where it has developed into a major pest in the production of crops (Goergen et al. 2016; Swamy et al. 2018; Lee et al. 2020; Sun et al. 2021). At present, FAW has been reported in more than 68 Eastern hemisphere countries (https://gd.eppo.int/taxon/LAPHFR/distribution) and colonisations have occurred regularly in low-latitude areas, causing great losses to agricultural production and disturbing the ecological balance (Day et al. 2017; Jing et al. 2020). Moreover, FAW has a predatory habit and can attack other sympatric pests and some natural enemies, showing advantages in niche competition (Wu 2020; Song et al. 2021). Given the fact that FAW can spread rapidly into different geographical areas with varying climatic conditions in the Eastern hemisphere, reflecting its strong environmental adaptability, it has a considerable potential to invade other regions in the future. Therefore, the development of effective control strategies for FAW has become an urgent task and more research is currently needed to achieve this goal.

In-depth knowledge on FAW genetic diversity and colonization history can help to develop targeted prevention and control strategies. The initial detection of invasive FAW was on the island country São Tomé and Príncipe in April 2016, followed by outbreaks on mainland western Africa (Benin, Nigeria, Ghana) in June 2016 (Goergen et al. 2016; Cock et al. 2017; Nagoshi et al. 2019). Studies have suggested that FAW invaded Africa through international trade, although it has strong long-distance migration ability (Johnson 1987; Chapman et al. 2017; Cock et al. 2017). However, the specific origin(s) of the invasion is/are still unclear, and there are some different hypotheses based on different research methods. Studies based on molecular markers indicated that the invasive FAW originated from a single source with a high similarity to samples from Florida and the Greater Antilles (Nagoshi et al. 2017; Nagoshi et al. 2020), while other studies using genome-wide analyses supported multiple introductions of FAW into Africa and suggested South America as a potential source of FAW invasion into Africa (Nagoshi et al. 2022; Tay et al. 2022). In the Western hemisphere, the FAW has undergone a divergence into two “strains”, a rice-strain and a corn-strain accordingly (Pashley et al. 1985; Pashley 1986; Pashley et al. 1992). The former is thought to preferentially feed on rice and various pasture grasses and the latter on maize and sorghum (Nagoshi and Meagher 2004; Prowell et al. 2004; Machado et al. 2008), however, this pattern is not consistent (Juárez et al. 2012; Murúa et al. 2015). Although the two strains exhibit allochronic differentiation in their mating time and sex pheromone composition (Pashley et al. 1992; Groot et al. 2008; Schöfl et al. 2009), there is no absolute reproductive isolation between them (Nagoshi et al. 2006; Groot et al. 2010). The strains are morphologically identical but can be distinguished using molecular markers such as the mitochondrial cytochrome oxidase I (*COI*) and the Z-chromosomal triosephosphate isomerase (*Tpi)* gene fragments (Nagoshi et al. 2007; Nagoshi 2010; Nagoshi 2012). Previous studies based on molecular markers and genome-wide analysis showed that the invasive samples in the Eastern hemisphere were inter-strain hybrids with a dominant corn-strain background (Nagoshi et al. 2019; Zhang et al. 2020). But a divergent *Tpi* haplotype in the Eastern hemisphere was detected with 10 unique single nucleotide polymorphisms (SNPs) compared to previously reported corn-or rice-strain genotype in the Americas, suggesting a more complex genetic origin of invasive populations (Nagoshi 2019; Zhang et al. 2020). In addition, the invasive FAW has also been reported in China to feed on rice and various weeds in the field (Kalleshwaraswamy et al. 2019), and it has also been confirmed that the invasive colony can complete its life cycle on rice and various grasses in the laboratory (Su et al. 2022). Therefore, the specific strain identification and genetic origin of the invasive FAW needs to be confirmed.

In recent years, intensive research has been undertaken on the biology, insecticide resistance, and management of FAW populations (Gichuhi et al. 2020; Jiang et al. 2021; Shan et al. 2022; Withers et al. 2022). However, our knowledge of the differentiation of the two strains are yet not fully understood, although there have been many comparative studies based on genome and transcriptome data focused on gene copy numbers and detoxification related genes (Gimenez et al. 2020; Nam et al. 2020; Xiao et al. 2020). In addition, the mechanisms underpinning the rapid spread of FAW in the Eastern hemisphere lacks in-depth understanding. In the Americas, studies have shown that several FAW populations have evolved resistance against a number of chemical pesticides and *Bacillus thuringiensis* (Bt) crops (Carvalho et al. 2013; Yang et al. 2018; Boaventura et al. 2020). Furthermore, the detoxification metabolism-related genes of the invasive populations in the Eastern hemisphere also displays a high level of variation (Guan et al. 2020; Zhang et al. 2020). Therefore, conducting a genome-level comparative study of FAW on a global scale will help to deepen our understanding of the evolution and rapid adaptation of this species.

In this study, we reveal the genetic structure and variation of global FAW populations and identify the most likely genetic origin of the invasive FAW populations, using a world-wide sample collection and whole-genome sequencing. Further, we analyze genomic signatures of selection to address the underlying mechanism associated with adaptive diversity and population evolution, which provides a solid basis for the early warning and prevention of FAW, and a genomic framework for future research on this invasive pest.

## Results

### Global genetic structure of FAW populations

We sampled 153 FAW individuals of invasive populations from the Eastern hemisphere (113 in Asia and 40 in Africa) for Illumina sequencing (20× depth of coverage), and then jointly analyzed with 127 publicly available genomes of which 119 represent Western hemisphere samples (Supplementary Table 1). In total, clean reads of 280 samples were individually aligned to the reference genome (390.38-Mb chromosome-level FAW genome, GenBank accession number: WUTJ00000000.2) with genome coverage ranging from 71.22% to 94.99% (Supplementary Table 2). A dataset of 39,158,713 SNPs was obtained after preliminary filtering, and then a total of 5,124,500 high-quality SNPs were finally retained for further analyses.

Using the high-quality SNP data, a neighbor-joining phylogeny of 280 FAW samples was constructed with the congeneric *Spodoptera litura* as an outgroup (Fig. 1a). Clustering results revealed that the invasive samples from the Eastern hemisphere formed a large cluster nested within the Western hemisphere samples. In the whole Eastern hemisphere branch, samples from Africa and Asia are mixed in different clusters, harbouring less genetic variation than those from the Americas. The clusters did not correlate with sampling time or geographic origin. For the Western hemisphere branch, several clusters were formed with high bootstrap values and most samples from the same laboratory strain were clustered together. As in the Eastern hemisphere, these clusters in the Western hemisphere did not group by geographic origin.

**Fig. 1.**
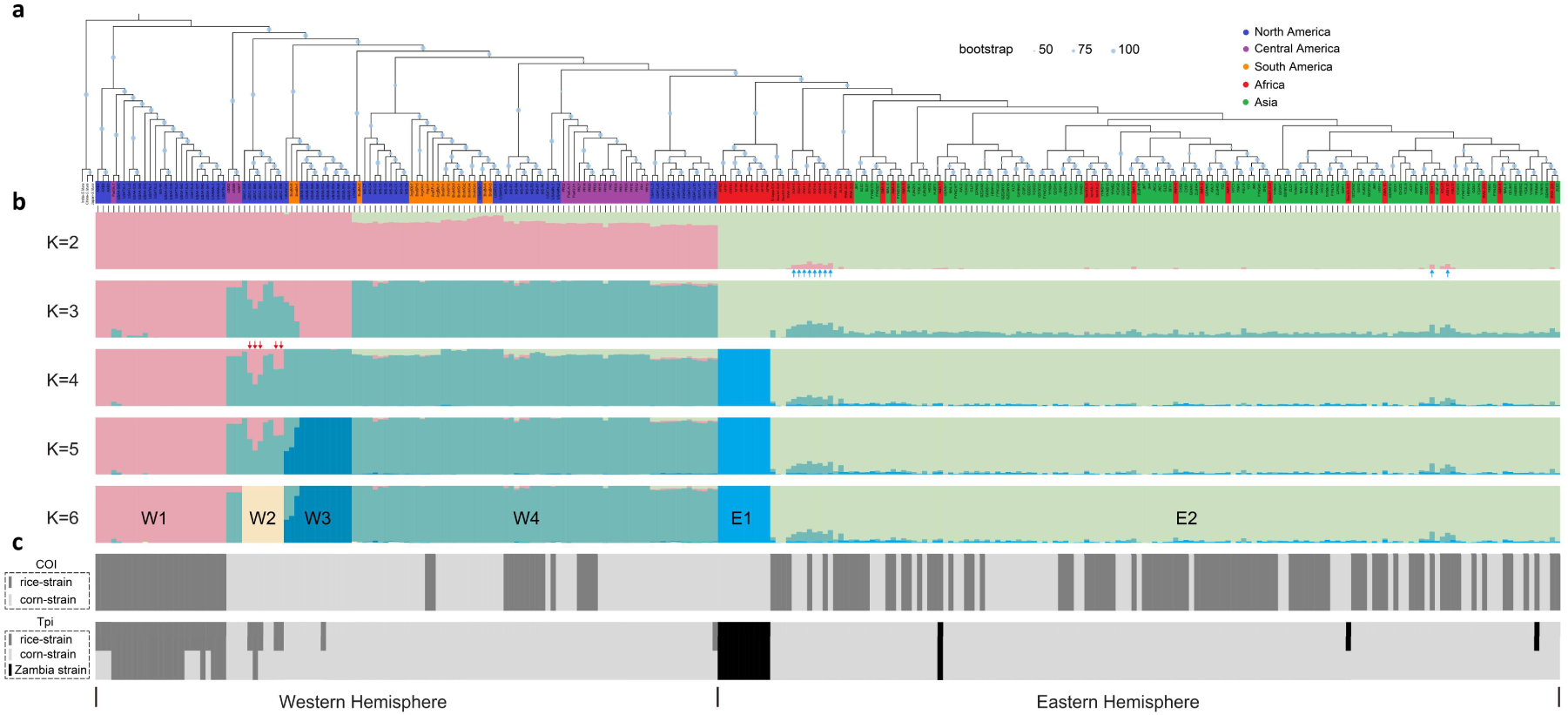
Strain identification and population structure of 280 FAW samples. **(a)** Neighbor joining (NJ) clustering of FAW samples, three *Spodoptera litura* samples were used as outgroup, colors for each sample ID represent different geographic regions, corresponding to Asia (green), Africa (red), North America (blue), Central America (purple), South America (orange). **(b)** Population structure of FAW samples by ADMIXTURE, the K value shows the assumed different number of ancestral populations, K=6 is the best fitting admixture model suggested by the cross-validation error. W1-W4 represent four populations in the Western hemisphere. E1-E2 represent the two invasive populations in the Eastern hemisphere. Blue arrows indicate ten samples from Ghana, red arrows indicate five samples corresponding to the five samples fall in W1 circle in Fig. 2c. **(c)** Strain identification based on the full-length mitochondrial *CO*I gene and the Z-chromosomal *Tpi* gene. Light grey indicates the corn-strain, dark grey indicates the rice-strain, black indicates the Zambia strain. Samples containing two colors represent a heterozygous haplotype.

Principal component analysis (PCA) and ADMIXTURE analysis were performed to further analyze the genetic structure of the FAW populations. Consistent with the cluster analysis, the ADMIXTURE results clearly distinguish the samples from the Eastern and Western hemispheres at K=2, where K is the assumed number of ancestral populations (Fig. 1b). When we increased the K value to K=4, the Western hemisphere samples were divided into two main populations, and the Eastern hemisphere samples were also divided into two populations (Supplementary Fig. 1, Supplementary Fig. 2), in which the E1 population represents a laboratory strain originating from field samples collected from Zambia in Africa in 2017 (Zambia strain), while the E2 population represents the remaining invasive samples collected in field. This result, of two distinct populations in the Eastern hemisphere, were also supported by the PCA, in which the first two PCs explain 6.26% and 4.01% of the total variation and separate the Zambia strain from the rest of the Eastern hemisphere samples (Fig. 2b). The cross-validation error suggested that K=6 was the optimal model of ADMIXTURE, resulting in four divergent Western hemisphere populations (Fig. 1b, Supplementary Fig. 3). The W1 population contains 24 North American samples and one Central American sample, which corresponds to the genetically most divergent cluster in the distance tree. All the 25 samples in this population showed a rice-strain genotype at both the mitochondrial *CO*I and the Z-chromosomal *Tpi* genes, with six samples exhibiting a heterozygous *Tpi* genotype (Fig. 1c). This population also included the previously characterized rice strain (Gouin et al. 2017) samples (AXE3, AXE5, AXE6) and 10 field collected (Gui et al. 2022) samples (USAA) feeding on broadleaf signalgrass, *Urochloa platyphylla*, often considered to be a rice-strain host. The W4 population contains 70 samples from North-, Central-, and South America, including three previously characterized corn strain (Gouin et al. 2017) samples (ASW2, ASW6, ASW7), which could be considered as representative American corn-strain population (Fig. 1c). The W2 and W3 populations mainly correspond to two previously reported laboratory strains (Gui et al. 2022). It is worth noting that some samples in the W2 population showed a heterozygous *Tpi* genotype combining a rice-strain allele with a corn-strain allele, which indicated that the W2 population might be an admixed group (Fig. 1c). This was also reflected in the PCA results, in which W2 samples grouped into the W1 or W4 clusters according to the *Tpi* genotype (Fig. 2c).

**Fig. 2.**
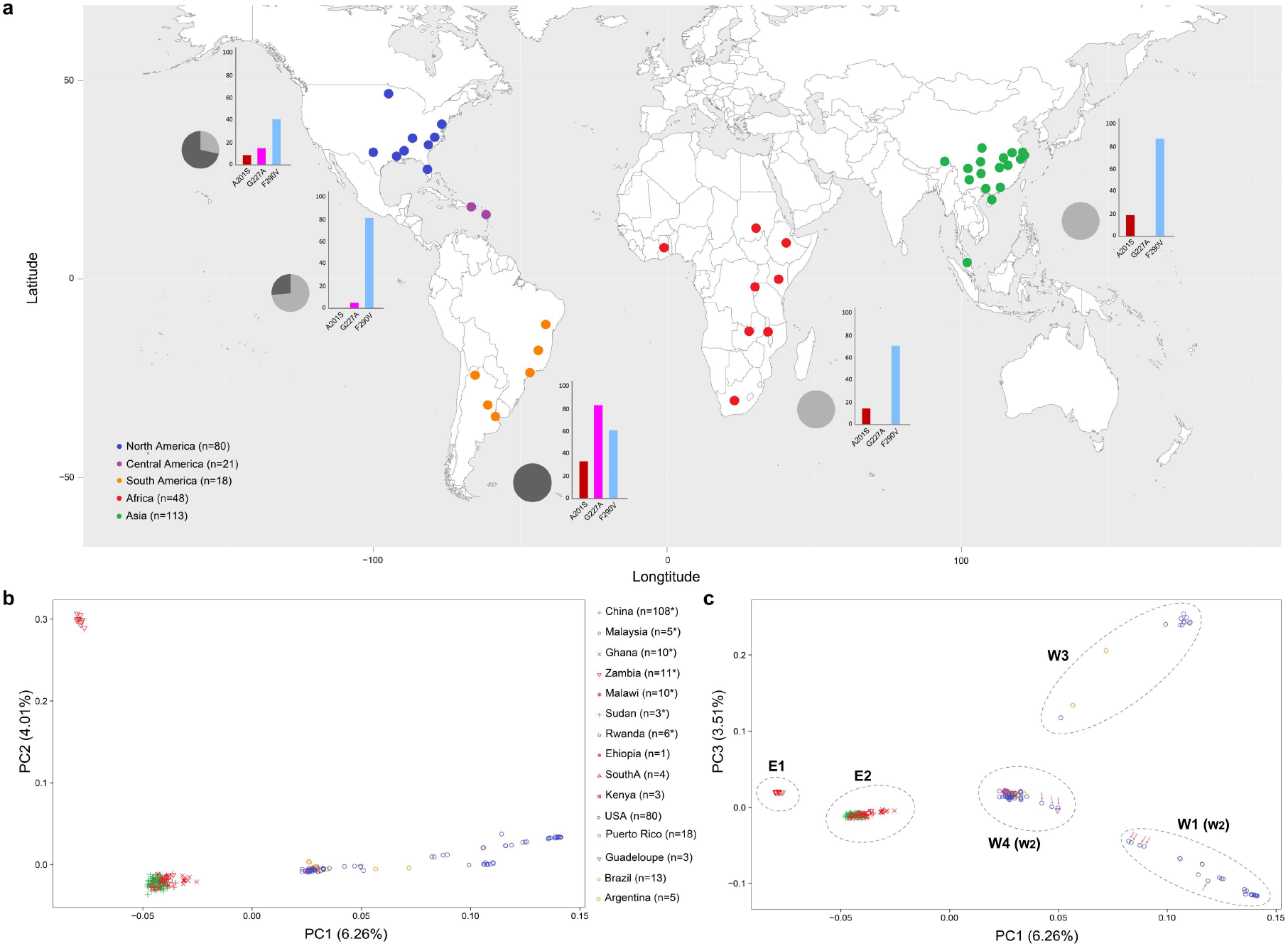
Sample locations and genetic structure of 280 FAW samples. **(a)** Overview of sample locations showing the invasive samples collected from the Eastern hemisphere (green and red dots) and the original samples from the Western hemisphere (blue, purple and orange dots). The pie chart represents the proportion of Texas-type (dark grey) and Florida-type (light grey) in mitochondrial corn-strain FAW samples. The histogram represents the proportion of the three resistance variant sites of *AChE* gene in FAW samples from different continents. **(b)** PCA plot of the first two components using whole-genome SNPs. Number of samples per country are indicated, numbers with an asterisk represent samples newly sequenced in this study. **(c)** PCA plot of the first and the third components, the dotted circles correspond to the clustering results in ADMIXTURE when K=6, red arrows indicate eight samples from the W2 population, in which five samples fall in the W1 circle corresponding to the five sample with red arrows in Fig. 1b when K=4. All samples within the W1 (W2) circle contain at least one rice-strain *Tpi* allele.

In summary, the global genetic structure of FAW populations based on distance clustering, PCA as well as ADMIXTURE analysis showed congruent results. The FAW population structure was not related to geographical origin in either the Eastern or Western hemisphere, while the invasive Eastern hemisphere samples formed a distinct cluster from the Western hemisphere samples. The Western hemisphere populations showed a high degree of differentiation, which is strongly associated with Z-chromosomal *Tpi* gene haplotype, which was previously used for strain identification in the Americas (Fig. 1). The invasive field samples in the Eastern hemisphere shared similar genetic background, which is consistent with a common origin and a recent invasion. In addition, the Zambia strain that may represent a relatively differentiated population was detected in the Eastern hemisphere.

### Mitochondrial haplotype network

The complete mitochondrial genome sequences of 240 samples were successfully assembled, including 134 samples from the Eastern hemisphere and 106 samples from the Western hemisphere. A conserved sequence set of these 240 samples, with 14930 bp in length, which contained all the mitochondrial protein-coding genes, tRNAs and rRNAs, was used to perform a haplotype network analysis. A total of 78 haplotypes were identified (Fig. 3, Supplementary Table 3). The vast majority of these (70 haplotypes; 65.1%) were identified in the Western hemisphere samples; consistent with it being the native range. Only nine haplotypes (6.7%) were identified in the invasive Eastern hemisphere samples, suggesting multiple colonizing individuals. Only one haplotype (Hap3) was shared by both Western and Eastern hemisphere samples.

**Fig. 3.**
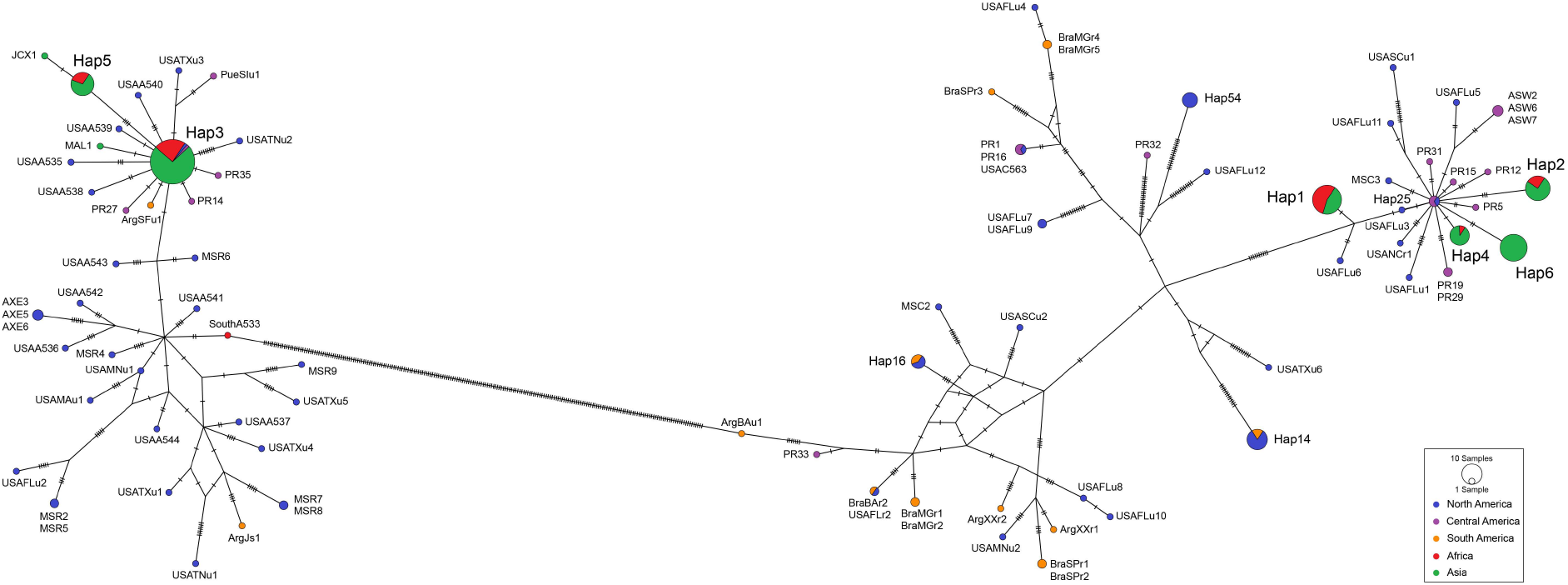
Haplotype networks of 240 FAW complete mitochondrial genomes. Different colors represent geographic regions consistent with Fig. 1 and Fig. 2. The left part indicates the rice-strain haplogroup based on mitochondrial *CO*I genotype, the central Hap3 consists of mostly invasive samples and those from Central-and North America. The right part indicates the corn-strain haplogroup, invasive haplotypes were evolved from the central Hap25, which consists of samples from Central-and North America, samples from Zambia strain grouped in the divergent Hap1 in corn-strain haplogroup.

Through haplotype network analysis, all 240 FAW samples were clearly divided into a “corn-strain” haplogroup and a “rice-strain” haplogroup, indicating that the two haplogroups have experienced deep mitochondrial genome divergence (Fig. 3). The corn-strain and rice-strain haplogroups contain 42 and 36 haplotypes, respectively, with the common feature that both haplogroups do not show any structure associated with geographic origin (Fig. 3). The Western hemisphere samples represented the largest variation across both corn-and rice-strain haplogroups whereas invasive samples from the Eastern hemisphere contained only nine haplotypes with relatively small differences between haplotypes within each haplogroup. In the rice-strain haplogroup, all invasive samples but one, from South Africa (SouthA533), were grouped in the cluster with Hap3 as the central haplotype, in which one sample each from North America (USAMAu2) and Central America (PR30) were included. The other major invasive haplotype, Hap5, consisting of 14 invasive samples, showed four variation sites compared to Hap3. In the corn-strain haplogroup, invasive samples were distributed across four haplotypes that formed a cluster with Hap25 as central haplotype, which consists of one sample from North America (MSC7) and two samples from Central America (PR18, PueGUr1). Notably, Hap6 only contained 19 Asian samples with two variation sites compared to the Hap25 haplotype. In addition, the Hap1 haplotype consists of 10 laboratory samples (Zambia strain) and 12 field samples from Asia and Africa, share a unique haplotype at least four nucleotide differences away from the other three corn-strain invasive haplotypes Hap2, Hap4, and Hap6.

In general, the mitochondrial haplotype network showed no correlation with geographic origin, nor with nuclear genome-wide population structure. However, invasive haplotypes are often associated with those from Central-and North America. It is obvious that mitochondrial genome differentiation in the invasive Eastern hemisphere samples was significantly lower than in Western hemisphere samples of either rice-strain or corn-strain genotype. Samples from Asia always shared haplotypes with those from Africa, except Hap6, which is until now only detected in China. By comparing the mitochondrial genome sequences, the differences between Eastern hemisphere haplotypes contain length variation caused by the simple sequence repeats located in protein-coding gene (NADH dehydrogenase 6) and tRNA genes (tRNA-Asp, tRNA-Glu) (Supplementary Fig. 4), while sequence differences in Western hemisphere samples are mainly point mutations.

### The genetic origin of invasive FAW

Results based on distance tree, PCA and ADMIXTURE analysis (K=2) distinguished the samples from the Eastern and Western hemispheres, indicating that the invasive samples are genetically different from those in the Western hemisphere. Moreover, the invasive individuals can be further divided into two well-differentiated populations based on PC2 and the ADMIXTURE result when K=4. Therefore, we performed further comparison of samples from the Eastern and Western hemispheres.

First, we calculated the heterozygosity of samples from different FAW populations. The results showed that samples from W1 population had the highest average heterozygosity (Fig. 4a). However, samples from the above region varied in heterozygosity. The average heterozygosity of invasive samples (E2 population) were also at relatively high levels, with all samples had a similar heterozygosity. Second, we calculated the nucleotide diversity (Pi) of the six populations, among which the three laboratory populations (E1, W2, W3) had relatively low nucleotide diversity (Fig. 4b). Unexpectedly, the Pi value of the invasive field population (E2) is higher than that of the W1 and W4 populations which represent the two highly differentiated populations in Americas, indicating that the E2 population has a highly admixed background. Finally, by calculating the genetic differentiation (Fst) between six populations, the E2 population showed the lowest Fst with the American corn-strain population W4 (Fst=0.06, Supplementary Table 4), indicating that these two populations have a closer genetic relationship. The *Tpi* genotype of samples from the E2 population exclusively showed the corn-strain or Zambia-strain haplotype (Fig. 1c). Given that all nuclear data suggest an origin from W4 corn-strain, we assume that the rice-strain mitochondrial haplotypes in the Eastern hemisphere originated from W4 corn-strain with rice-strain mitochondrial haplotype.

**Fig. 4.**
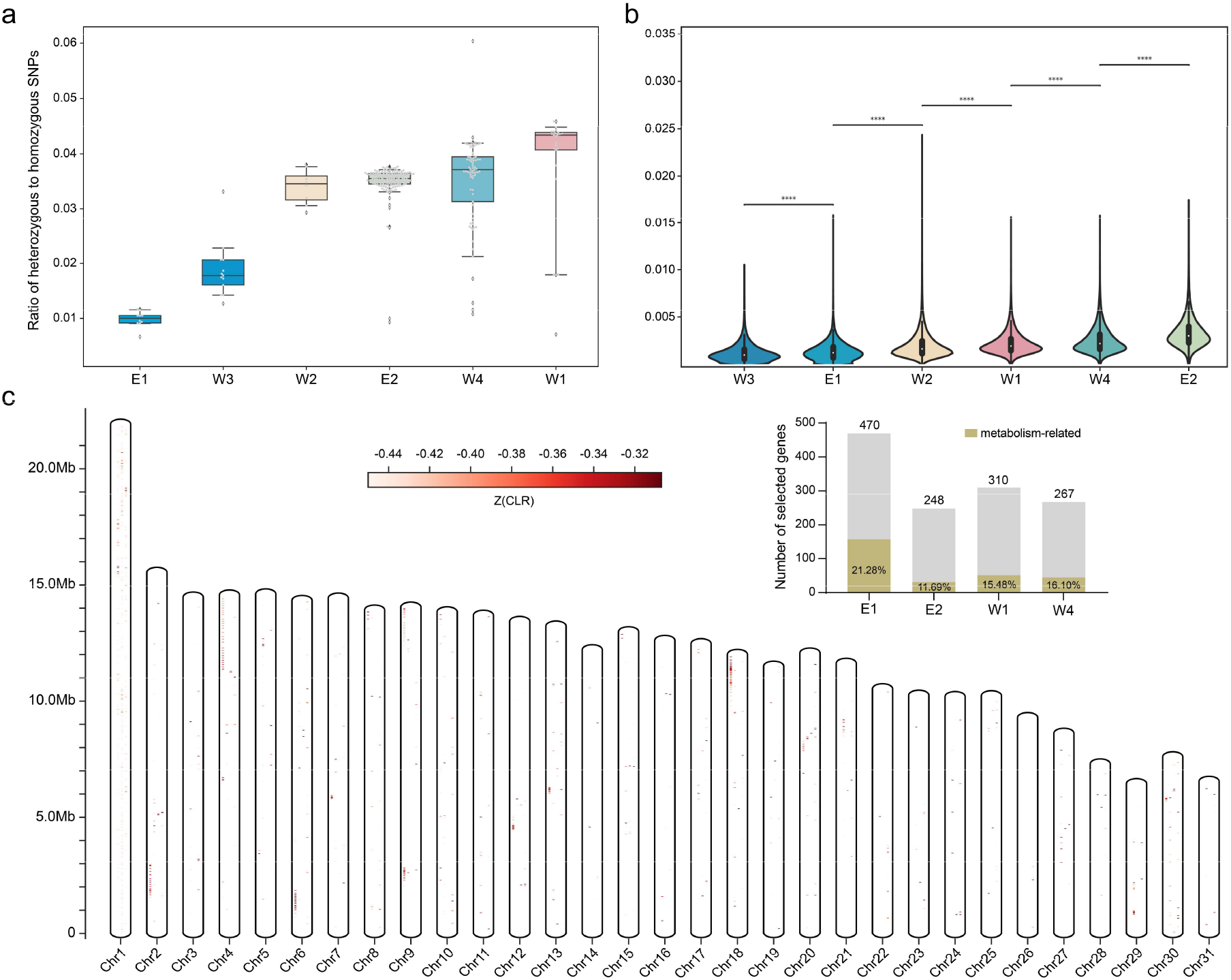
Genetic diversity and selective sweeps of different FAW populations. **(a)** Ratio of heterozygous to homozygous SNPs in samples from six FAW populations. **(b)** Genome-wide nucleotide diversity (Pi) of six FAW populations. **(c)** Signatures of selective sweep for four different FAW populations, the four columns in each chromosome represent the E1, E2, W1, W4 populations from left to right. The bar graph in the upper right corner indicates the number of candidate genes under selection for each population. Numbers inside the colored bars represent the proportion of metabolism-related genes.

It is worth noting that there is a relatively independent E1 population in the Eastern hemisphere, an established laboratory strain originating from Zambian field samples (Zambia strain). Both PCA and ADMIXTURE results showed that this population was significantly separated from the invasive field population (Fig. 1b, 2b, 2c). We previously assembled the genome of this Zambia strain and identified its unique *Tpi* haplotype and mitochondrial insertion into nuclear genome (Zhang et al. 2020). Here, we scanned genomic data of all 280 samples in this study, the results showed that the Zambia strain *Tpi* haplotype was only distributed in the Eastern hemisphere with a proportion of 2% in field individuals (Fig. 1c, Supplementary Table 1). The mitochondrial haplotype of this population (E1) was located in Hap1, which also represented a unique haplotype different from the main invasive haplotypes in the corn-strain branch (Fig. 3). These results, based on whole genomic comparison, mitochondrial haplotype and *Tpi* haplotype, reinforce the possibility that the Zambia strain may represent a different genetic background distinct from the known corn-strain or rice-strain of American origin. In addition, mitochondrial haplotype of E1 population (Hap1) was detected in 12 samples from E2 population, and three samples with Zambia-strain *Tpi* haplotype were detected in Asian and African field samples. The above results indicate that there is genetic introgression of this Zambia strain (E1) into the invasive field population (E2), which was also supported by the ADMIXTURE analysis in which small but noticeable E1 components were detected in the E2 population (Fig. 1b). Based on the above analysis, the invasive field individuals with high heterozygosity and nucleotide diversity in the Eastern hemisphere could be explained by the mixing of two different backgrounds. Our results tend to support the hypothesis that invasive FAW was derived from the hybrid offspring of the Zambia strain (E1) and multiple genotypes from within the American corn-strain (W4).

### Selective sweep and transcriptome analysis

In order to clarify the differentiation of different FAW populations, we conducted selective sweep analysis using the composite likelihood ratio (CLR) method. The genetic backgrounds of W2 and W3 populations are not stable based on PCA results and strain identification using the *Tpi* gene (Fig. 1c, 2c). Therefore, the remaining four populations were selected for CLR analysis (Fig. 4c), including two representative early differentiated populations namely American rice-strain (W1) and corn-strain (W4) in the Western hemisphere, and two populations namely Zambia strain (E1) and invasive filed population (E2) in the Eastern hemisphere. The genes detected by CLR methods for each population were considered as candidates for selective sweeps (Fig. 4c). The results showed that the E1 population had the most genomic selection signals, resulting in 470 selected genes of which 21.28% were relevant to metabolic pathways such as the ABC transporter family and cytochrome P450 genes. 248 selected genes were identified in the E2 population with a proportion of 11.69% relevant to metabolic pathways, which is much lower than that of the E1 population. 310 and 267 genes under selection were identified for the American rice-strain W1 and corn-strain W4 populations, respectively. However, the KEGG enrichment analysis of these selected genes did not show significant differences between the two host-plant populations (Supplementary Fig. 5), and the two populations have similar proportions of metabolism-related genes under selection (Fig. 4c).

To further analyze the feeding preference of the invasive FAW in the Eastern hemisphere, two representative laboratory strains corresponding to the Zambia strain (E1 strain) and an invasive field population (E2 strain) were reared on either maize or rice and sampled for comparative transcriptome analysis. The results showed that the highest number of differentially expressed genes was observed at 48 hours for both strains, when feeding on plants compared to artificial diet (Fig. 5). Surprisingly, neither strain showed significant differences in feeding on different host plants at 48 hours, with a similar number of differentially expressed genes (DEGs) for either maize or rice (Fig. 5). However, it is obvious that the main difference was between strains, regardless of the host plants, the E1 strain had more DEGs than the E2 strain, indicating that the E1 strain had experienced a stronger reaction when feeding on these two host plants. We further analyzed the up-regulated DEGs of the two strains at 48 hours. The KEGG enrichment analysis showed that the DEGs in the E1 strain were mainly enriched in the metabolic pathway such as drug metabolism, while the DEGs in E2 strain were mainly enriched in the immune pathway such as toll and Imd signaling pathways (Supplementary Fig. 6, 7). The above results showed the obvious difference in feeding response between two strains, which again indirectly reflect the different genetic background of E1 and E2 populations.

**Fig. 5.**
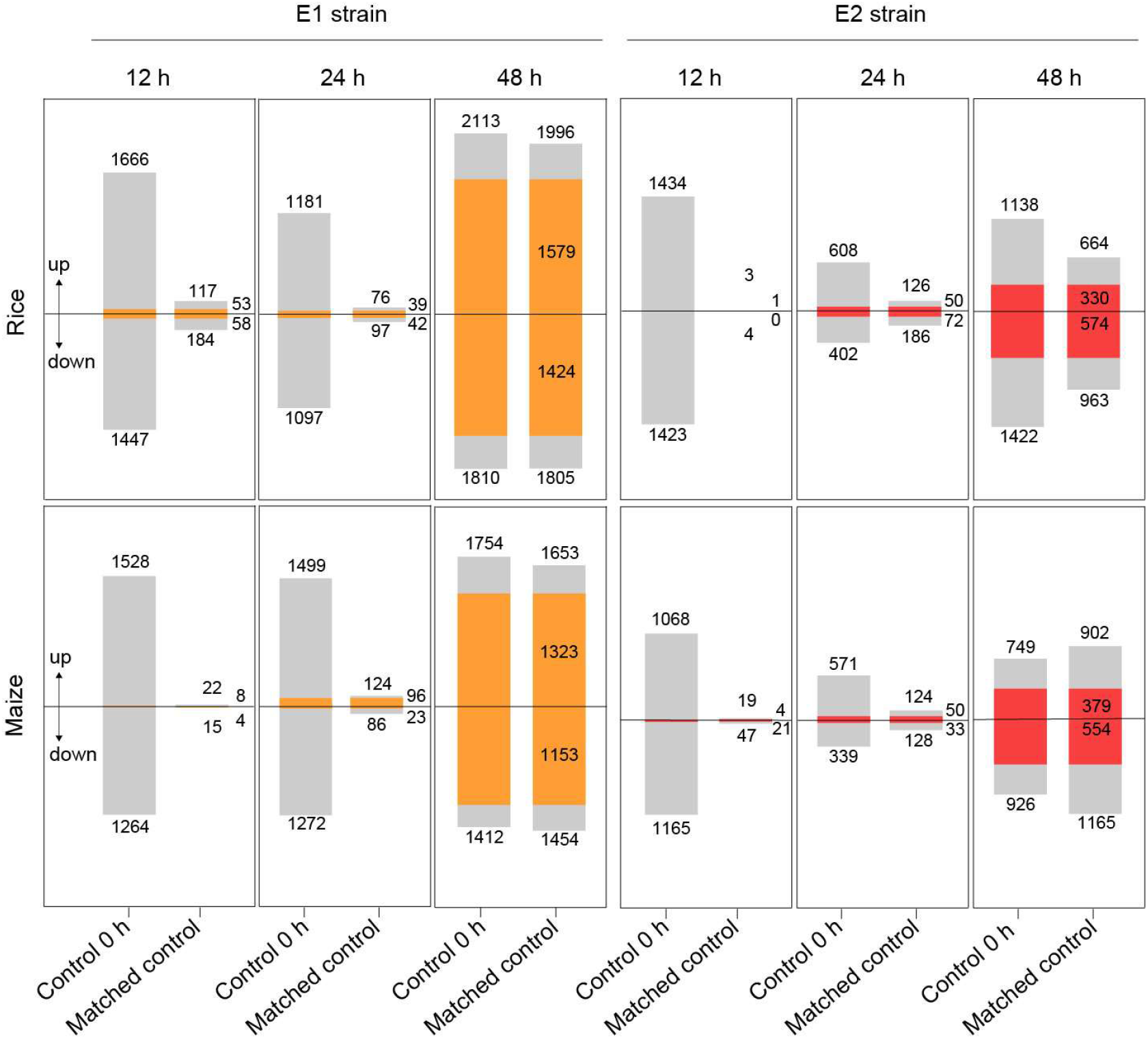
Identification of DEGs of two invasive strains in response to different host plants. Number of DEGs between two strains feeding on maize and rice. For each time point, “control 0 h” means the number of DEGs relative to the control sample collected at time 0; “matched control” means the number of DEGs relative to the control sample collected at the matching time point (12 h, 24 h, 48 h). Numbers inside the colored bars represent DEGs in both comparisons.

### Genome-wide scan of insecticide resistance genes

According to the previously reported resistance mutations of *S. frugiperda*, including Bt-related ATP-dependent Binding Cassette subfamily C2 (*ABCC2*) gene (Boaventura et al. 2020) and pesticide-related genes (Carvalho et al. 2013; Boaventura et al. 2020) such as acetylcholinesterase (*AChE*), voltage-gated sodium channel (*VGSC*), ryanodine receptor (*RyR*), we carried out genome scan of these four genes in all 280 FAW samples. The result showed that all four resistance-related genes exhibited high levels of nucleotide substitutions with an overall ratio between 15.76% - 42.78% in the coding sequence (CDS) region. The non-synonymous mutation ratio at amino acid level varied from 3.40% to 9.48% (Supplementary Table 5), with the mutation rates of *VGSC* and *RyR* genes significantly lower than those of *ABCC2* and *AChE* genes, indicating that the latter two genes were experiencing greater selection pressure. However, only three known resistance mutations in *AChE* gene were detected, which were related to organophosphate insecticide resistance. For the three resistance mutation sites (A201S, G227A, F290V), the distribution frequency varied among samples from different geographic regions (Fig 2a, Table 1). The G227A mutation was not detected in any Eastern hemisphere samples, but was widely distributed in the Western hemisphere, especially in South America where the prevalence reached 83.3%. Interestingly, no homozygous G227A mutant was detected in any of the 280 FAW samples in this study, which implies that this could be a critical locus associated with lethality.

**Table 1.** Statistics of resistance mutation sites of the *AChE* gene in FAW samples from different continents.

| *AChE* | A201S | G227A | F290V |
| --- | --- | --- | --- |
| Asia | 18,58% | 0 | 86,72% |
| Africa | 14,58% | 0 | 70,83% |
| North America | 8,75% | 15% | 40% |
| Central America | 0 | 4,76% | 80,95% |
| South America | 33,33% | 83,33% | 61,11% |

## Discussion

The current study covers the most extensive analysis of FAW genomic data so far and includes the distribution range of FAW in the region of origin and invasive areas. Although Asian samples were mainly from China, previous results showed that samples from Southeast Asia and Australia shared a similar genetic background (Nagoshi et al. 2020; Tay et al. 2022). In contrast to previous studies suggesting one panmictic population of American samples based on whole genome comparisons (Schlum et al. 2021), the global FAW population structure based on genome-wide SNPs in this study showed that there were several clearly differentiated populations in the Western hemisphere but with little correlation to geographical distribution, which could be caused by the frequently migratory behavior and crop transport of FAW. In general, the population structure in the Western hemisphere showed a correlation with the Z-chromosomal *Tpi* gene, but not with the mitochondrial *CO*I gene (Fig. 1b, 1c). However, all samples in the identified W1 population showed a consistent rice-strain genotype for both mitochondrial *CO*I and *Tpi* genes, indicating that there may be a certain degree of reproductive isolation or a one-direction mating preference between rice females and corn males, which was also supported by a previous study regarding the existence of post-zygotic incompatibilities between the corn-and rice strain (Pashley and Martin 1987; Dumas 2015). Then after differentiation, two distinct host strains mixed but still conserved their main nuclear genomic information to a certain extent. Supporting this assertion, rice-strain samples from diverse locations of America collected in different years clustered together and showed a similar genetic background based on the results of genome-level analysis (Fig. 2c, Supplementary Fig. 2).

Revealing the origin has always been a key issue in the study of biological invasions. The present study indicates that the invasive field FAW in the Eastern hemisphere showed the closest similarity to the American corn-strain population (W4) from the Western hemisphere, and the *Tpi* genotype was also identical to the American corn-strain except for a few samples containing the Zambia-strain *Tpi* haplotype (Fig. 1c). However, the W4 population identified in this study contains samples from North America, Central America, and South America, so the specific origin cannot be determined yet with certainty. Through haplotype network analysis, mitochondrial rice-strain Eastern hemisphere samples share a haplotype (Hap3) with samples from North-and Central America. The corn-strain haplotypes of the invasive samples were also similar to the Western hemisphere haplotype (Hap25) consisting of North-and Central America samples. This indicates that the invasive samples might be more closely related to the North-and Central American samples at the mitochondrial genome-wide level, although this could also reflect a lack of sampling in South America. According to previous studies, mitochondrial corn-strain FAW could be divided into Texas (TX) and Florida (FL) types according to the frequency of two variable sites in the *CO*I gene (Nagoshi et al. 2007; Nagoshi et al. 2008). The TX-type was distributed throughout the Western hemisphere, while the FL-type was limited to the eastern coast of the United States, extending southward to Central America to areas such as Puerto Rico and the Lesser Antilles (Nagoshi et al. 2010; Nagoshi et al. 2015). The results of our study showed that all corn-strain samples in the Eastern hemisphere were FL-types (Fig. 2a), which also indicated that the invasive FAW may originate from North-and Central America. In addition, the resistance-associated locus (G227A) in the *AChE* gene had a mutation frequency of 83.3% in South America samples, but was not detected in the Eastern hemisphere (Fig. 2a), again suggesting that the invasive FAW was less likely to be derived from South America.

For the Zambia strain (E1 population) identified in the Eastern hemisphere, the results based on whole genome data clearly isolated this population from the rest of the invasive population samples. Previous population clustering based on microsatellites also determined a weak genetic structure between invasive FAW populations in Africa, in which FAW from Zambia were more genetically isolated from other countries (Withers et al. 2021). Unfortunately, we did not find any individuals with the E1 genotype at whole-genome level in field-collected samples in our study. Giving that Zambia strain (E1 population) is a laboratory strain reared for about 12 generations before sequencing, there is bound to be some selection and genetic drift during two years of rearing in the laboratory. However, previous studies showed small genetic differentiation between laboratory stains and field-collected groups for American FAW samples (Schlum et al. 2021). Besides, there were also several laboratory strains in this study that did not form an independent population, such as AXE3, AXE5, AXE6 reared in laboratory for more than 15 years (Gouin et al. 2022) nested within the W1 population. The most distinctive feature of the Zambia strain is the divergent *Tpi* gene haplotype, which has so far never been detected in the Western hemisphere (Nagoshi 2019; Nagoshi et al. 2019; Zhang et al. 2020). This divergent haplotype was also detected in some Asian (RFAW4) and African (SouthA-523, AMA7) field samples (Supplementary Table 1). In Africa, there are many *Spodoptera* species, which are closely related to *S. frugiperda*. We therefore assessed whether the Zambia strain might be an interspecific hybrid. However, based on molecular identification of 28s rDNA (28S) and elongation factor-1a (ef1a) nuclear markers (Kergoat et al. 2012), as well as the result of high mapping rate and coverage to the reference genome, all results indicated that Zambia strain was *S. frugiperda* and might represent a more divergent lineage. Based on this, we speculate that the Zambia strain may derive from an independently invaded FAW population in Africa. Subsequently, the Zambia strain hybridized with the recently (in 2016) introduced American corn-strain population, after which the population expanded leading to the outbreak in Africa. It is worth noting that the first detection of FAW was in São Tomé and Príncipe (Nagoshi et al. 2019), a small island country off the western equatorial coast of Central Africa, which could provide the basis for an independent evolution in a small-scale population. Moreover, Ghana and its surrounding areas in West Africa are likely to be the early invasive areas (Cock et al. 2017) for secondary contact, since the ADMIXTURE results indicated that the Ghana population (AGH1-10) contained a higher proportion of the American corn-strain genetic components (Fig. 1b).

It is important to clarify the genetic characteristics of the invasive FAW populations by comparative studies, which can also help to predict its feeding preference and response to control measures. Interestingly, the Zambia strain (E1) and the field strain (E2) in this study did not show significant differences in the transcriptome levels by feeding on rice or maize, and both strains could complete the life cycle normally. However, a large proportion of up-regulated genes of Zambia strain feeding on rice and maize were mainly enriched in the metabolism pathway, suggesting that this strain may not have evolved on rice or maize, but on other host plants. The genetic background of the invasive FAW in Eastern hemisphere is mainly characterized as corn-strain, which is consistent with the field damage on corn-strain host plants such as maize and sorghum (Goergen et al. 2016; Jing et al. 2020). However, due to the introgression of the Zambia-strain background, the invasive FAW has unclear damage characteristics and is at risk to damage more host crops. In conclusion, continuous field monitoring and genome-level comparative analysis are still needed to further explain the genomic introgression, to reveal the population differentiation and adaptive evolution of FAW in the newly invaded agro-ecosystem and different natural environments.

## Materials and Methods

### Sample collection

In this study, 153 invasive FAW larvae and adults were collected from 2017 to 2021, comprising 113 Asian and 40 African samples. The Asian samples included 108 field samples collected from China and five field samples from Malaysia. The African samples consisted of 30 field samples collected from five countries including Ghana (10), Zambia (1), Malawi (10), Sudan (3), Rwanda (6), the remaining 10 samples were from a laboratory strain collected from a maize field in Zambia in 2017 and reared for 15 months with about 12 generations in lab before sequencing, in Lancaster, UK. Field samples were collected as larvae or adults and were subsequently stored at −80°C. Samples were later identified as *S. frugiperda* by morphological and *COI* molecular identification. Genomes of a total of 127 samples were downloaded from a previous study (Gouin et al. 2017; Schlum et al. 2021; Gui et al. 2022), including 80 samples from the United States, 21 samples from Central America (Guadeloupe (3), Puerto Rico (18)), 18 samples from South America (Brazil (13), Argentina (5)), and eight samples from Africa (Kenya (3), Ethiopia (1), South Africa (4)). The geographical distribution and specific information of the samples are shown in Fig. 2a and Supplementary Table 1.

### DNA re-sequencing and SNP calling

The genomic DNA (gDNA) of all the samples in this study was extracted by using the Qiagen Genomic DNA kit followed by purity assessment and quantification with a NanoDrop One UV-Vis spectrophotometer (Thermo Fisher Scientific) and Qubit 3.0 Fluorometer (Invitrogen), respectively. A total of 0.5 μg gDNA for each sample was used as input to construct a library with an insert size of 250-350 bp. The library was sequenced on the Illumina NovaSeq 6000 platform for 150-bp paired-end sequencing. A total of 20× average genome coverage sequencing data was generated for each sample.

The newly sequenced (153 samples) and downloaded (127 samples) raw reads of 280 samples were filtered by trimming the adapter and low-quality regions using fastp v0.23.0 (Chen et al. 2018). The filtered reads were then aligned to the FAW genome using BWA-0.7.17 (r1188) mem software (Li 2013). Here, the chromosomal level high-quality FAW genome was used as reference genome (Zhang et al. 2020). SAMtools (Li et al. 2019) was used to sort the mapped reads and Picard (https://sourceforge.net/projects/picard/) was used to further remove the PCR duplicates. Variants were called independently for each sample by GATK (McKenna et al. 2020) HaplotypeCaller models and then were joined in a VCF file. Then a VCF file of 39,158,713 SNPs was obtained with the filter parameters “QD < 2.0 || MQ < 40.0 || FS > 60.0 || SOR >3.0 || MQRankSum < −12.5 || ReadPosRankSum < −8.0”.

### Genetic diversity, population structure and phylogeny

Heterozygosity of each sample was calculated using PLINK software (Purcell et al. 2007) based on the above-mentioned 39,158,713 SNPs. A subset of high-quality 5,124,500 SNPs was further filtered with PLINK software under “--max-missing 0.8 --maf 0.05 -- biallelic-only”. The filtered high-quality SNPs were further used for principal component analysis (PCA), distance tree construction and population structure analysis. The PCA was conducted using PLINK software with the first three eigenvectors plotted. VCF2Dis (https://github.com/BGI-shenzhen/VCF2Dis) was used to calculate the genetic distance matrix. Then a distance tree using neighbor-joining method was constructed by using PHYLIP v3.695 (http://evolution.genetics.washington.edu/phylip.html) with 1000 bootstrap replicates. Three congeneric *Spodoptera litura* individuals were used as an outgroup species (GenBank accession number: SRR5132417, SRR5132423, SRR5132429). The downloaded sequencing reads of *S. litura* were aligned to the FAW reference genome as described above with default parameters. The population structure was analyzed by ADMIXTURE v1.3.0 (-C 0.01) (Alexander et al. 2009) with the pre-defined genetic clusters increased from K = 2 to K = 8. Cross-validation was used to identify the best fitting model of ADMIXTURE. The fixation statistic (Fst) and genetic diversity (Pi) were used to estimate the genetic divergence within and between populations by VCFtools v.0.1.12b (Li 2011) with 10-kb non-overlapping windows. P-values for each comparison were calculated using the two-tailed Mann-Whitney U test with Bonferroni corrections.

### Mitochondrial haplotype network

Using the published mitochondrial genome of *S. frugiperda* as a reference (GenBank accession number: KM362176.1), Novoplasty v2.5.6 software (Dierckxsens et al. 2017) was used to assemble the mitochondrial genomes of all samples in this study. A manual correction was used to obtain the complete mitochondrial genome of each sample. Subsequently, the mitochondrial genomes of all samples were compared and aligned using MEGAX software (Kumar et al. 2018). Three fragments in intergenic regions (ND4-ND4L, tRNA-Leu-16sRNA, 12sRNA-tRNA-Met) were trimmed due to the low assembly confidence and the richness in simple repeat units such as A+T, resulting in a common length of 14930 bp (including all 13 protein-coding genes, 22 tRNA genes and 2 rRNA genes) for mitochondrial haplotype network analysis. DnaSP6 (Librado and Rozas 2009) was used to detect haplotypes, and then PopART (Leigh and Bryant 2015) was used to construct and visualize the haplotype networks with the “TCS network” method.

### Strain identification based on *Tpi* gene

Taking the full-length American corn-strain *Tpi* gene (1234 bp) as a reference (Gouin et al. 2017), the filtered genomic reads of all 280 FAW samples in this study were aligned to the reference *Tpi* gene sequence, and then gene variation of each sample was analyzed. We selected four stable variation sites in the coding sequence region of the *Tpi* gene to distinguish the American rice-and corn-strain FAW based on the 10 previously reported variation sites (Nagoshi 2012). Another two unique SNPs that belongs to the Zambia strain *Tpi* haplotype were also selected and this resulted in a final combination of six variable sites for identification of *Tpi* haplotype (Supplementary Fig. 8, Supplementary Table 6).

### Genome scan of resistance related genes and selective sweeps

Based on the previously reported insecticide resistance genes in *S. frugiperda* (Carvalho et al. 2013; Boaventura et al. 2020), the amino acid substitutions in AChE (A201S, G227A, F290V), VGSC (T929I, L932F, L1014F) and RyR (I4790M, G4946E) resulted in resistance to organophosphate, pyrethroid and diamide insecticides, respectively, and a 2-bp insertion in the *ABCC2* gene is linked to resistance to the Bt toxin Cry1Fa (Boaventura et al. 2020). The full-length sequences of these four genes were used as a reference to detect the variation in these genes in the genomic data of 280 FAW samples. Finally, the global genotype distribution of four previously reported SNPs associated with pesticide- and Bt-resistance was generated, and the proportion of synonymous mutations versus non-synonymous mutations was also calculated. For four representative populations from Eastern and Western hemisphere. A selective sweep analysis was conducted using SweeD 3.1 (Pavlidis et al. 2013) with a 10-Kb grid size. CLR was calculated by comparing the selective sweep model to the neutral model. Positions with CLR value above the top 1% cutoff level were extracted. Genes including these positions were identified to be under selection.

### Transcriptome analysis

For the invasive FAW in the Eastern hemisphere, two laboratory strains representing the Zambia strain (E1 strain) and the field strain (E2 strain, which is established from individuals collected in a maize field in Yunnan Province, China in 2019) were selected to carry out feeding experiments on different host plants. Larvae were grown on modified artificial diet (Ward’s Stonefly Heliothis diet, Rochester, NY) upto the late second instar stage. Subsequently larvae were starved for 24 hours and 20 early third instar larvae were selected and added together into a plastic box with a surplus of either rice or maize leaves, or with artificial diet (control group). At 0, 12, 24, and 48 hours, three larvae were sampled per treatment and immediately placed into liquid nitrogen and stored at −80°C. The feeding experiments were performed as three replicates.

Total RNA of each individual larvae was extracted using the RNeasy Mini extraction kit (Qiagen) according to the manufacturer’s protocol. The RNA integrity and purity were measured using a NanoPhotometer spectrophotometer (Implen) and Qubit 2.0 Flurometer (Life Technologies), respectively. Total RNA per sample was used to make indexed cDNA libraries using the NEB Next Ultra RNA Library Prep Kit for Illumina. The libraries had insert sizes of 250-300 bp and were sequenced on the Illumina NovaSeq 6000 platform with a paired-end 150-bp strategy. The transcriptome reads were trimmed using Trimmomatic v0.38 (Bolger et al. 2014) and then mapped to reference FAW genome by using Hisat2 v2.1.0 (Kim et al. 2019) with default parameters. The relative abundance of genes was measured in FPKM using StringTie v2.1.1 (Pertea et al. 2015) and gene was excluded when the FPKM values < 0.1 in at least one sample. Then, differentially expressed genes (DEGs) were identified with DESeq2 (Love et al. 2014). The DEGs were determined with false discovery rate < 0.05, and an absolute value of log 2-fold change >1 between each pairwise comparison. To understand the function of DEGs, each DEG was aligned to Kyoto Encyclopedia of Genes and Genome (KEGG) database with E-value 10-5 and enrichment analysis were performed based on up-regulated DEGs.

## Supplementary materials

Supplementary figures and tables are available in the Supplementary material file.

## Data availability

All sequence data from this study have been deposited in GenBank under BioProject ID PRJNA591441.

## Acknowledgements

This work was supported by Shenzhen Natural Science Foundation (grant number JCYJ20200109150629266) and the Science and Technology Innovation Program of the Chinese Academy of Agricultural and Sciences.

## References

Alexander DH, Novembre J, Lange K. 2009. Fast model-based estimation of ancestry in unrelated individuals. Genome Res. 19: 1655–1664.

Boaventura D, Bolzan A, Padovez FE, Okuma DM, Omoto C, Nauen R. 2020. Detection of a ryanodine receptor target-site mutation in diamide insecticide resistant fall armyworm, Spodoptera frugiperda. Pest Manag Sci. 76: 47–54. https://doi.org/10.1002/ps.5505

Boaventura D, Ulrich J, Lueke B, Bolzan A, Okuma D, Gutbrod O, Geibel S, Zeng Q, Dourado PM, Martinelli S, et al. 2020. Molecular characterization of Cry1F resistance in fall armyworm, Spodoptera frugiperda from Brazil. Insect Biochem Mol Biol. 116: 103280. https://doi.org/10.1016/j.ibmb.2019.103280

Bolger AM, Lohse M, Usadel B. 2014. Trimmomatic: a flexible trimmer for Illumina sequence data. Bioinformatics 30: 2114–2120. https://doi.org/10.1093/bioinformatics/btu170

Carvalho RA, Omoto C, Field LM, Williamson MS, Bass C. 2013. Investigating the molecular mechanisms of organophosphate and pyrethroid resistance in the fall armyworm Spodoptera frugiperda. PLoS One 8: e62268. https://doi.org/10.1371/journal.pone.0062268

Chapman D, Purse BV, Roy HE, Bullock JM. 2017. Global trade networks determine the distribution of invasive non-native species. Global Ecol Biogeogr. 26: 907–917. https://doi.org/10.1111/geb.12599

Chen SF, Zhou YQ, Chen YR, Gu J. 2018. fastp: an ultra-fast all-in-one FASTQ preprocessor. Bioinformatics 17: 884–890. https://doi.org/10.1093/bioinformatics/bty560

Cock MJW, Beseh PK, Buddie AG, Cafá G, Crozier J. 2017. Molecular methods to detect Spodoptera frugiperda in Ghana, and implications for monitoring the spread of invasive species in developing countries. Sci Rep. 7: 4103. https://doi.org/10.1038/s41598-017-04238-y

Day R, Abrahams P, Bateman M, Beale T, Clottey V, Cock M, Colmenarez Y, Corniani N, Early R, Godwin J, et al. 2017. Fall armyworm: impacts and implications for Africa. Outlooks Pest Manage. 28: 196–201. https://doi.org/10.1564/v28_oct_02

Dierckxsens N, Mardulyn P, Smits G. 2017. NOVOPlasty: de novo assembly of organelle genomes from whole genome data. Nucleic Acids Res. 45: e18. https://doi.org/10.1093/nar/gkw955

Dumas P. 2015. Spodoptera frugiperda (Lepidoptera: Noctuidae) host-plant variants: two host strains or two distinct species?. Genetica 143: 305–316. https://doi.org/10.1007/s10709-015-9829-2

Gichuhi J, Sevgan S, Khamis F, Van den Berg J, du Plessis H, Ekesi S, Herren JK. 2020. Diversity of fall armyworm, Spodoptera frugiperda and their gut bacterial community in Kenya. PeerJ. 8: e8701.

Gimenez S, Abdelgaffar H, Le Goff G, Hilliou F, Blanco CA, Hänniger S, Bretaudeau A, Legeai F, Nègre N, Jurat-Fuentes JL, et al. 2020. Adaptation by copy number variation increases insecticide resistance in the fall armyworm. Commun Biol. 3: 664. https://doi.org/10.1038/s42003-020-01382-6

Goergen G, Kumar PL, Sankung SB, Togola A, Tamò M. 2016. First report of outbreaks of the fall armyworm Spodoptera frugiperda (J E Smith) (Lepidoptera, Noctuidae), a new alien invasive pest in west and central Africa. PLoS One 11: e0165632. https://doi.org/10.1371/journal.pone.0165632

Gouin A, Bretaudeau A, Nam K, Gimenez S, Aury JM, Duvic B, Hilliou F, Durand N, Montagné N, Darboux I, et al. 2017. Two genomes of highly polyphagous lepidopteran pests (Spodoptera frugiperda, Noctuidae) with different host-plant ranges. Sci Rep. 7: 11816. https://doi.org/10.1038/s41598-017-10461-4

Groot AT, Marr M, Heckel GD, Schöfl G. 2010. The roles and interactions of reproductive isolation mechanisms in fall armyworm (Lepidoptera: Noctuidae) host strains. Ecol Entomol. 35: 105–118. https://doi.org/10.1111/j.1365-2311.2009.01138.x

Groot AT, Marr M, Schöfl G, Lorenz S, Svatos A, Heckel DG. 2008. Host strain specific sex pheromone variation in Spodoptera frugiperda. Front Zool. 5: 20. https://doi.org/10.1186/1742-9994-5-20

Guan F, Zhang JP, Shen HW, Wang XL, Padovan A, Walsh TK, Tay WT, Gordon KHJ, James W, Czepak C, et al. 2020. Whole-genome sequencing to detect mutations associated with resistance to insecticides and Bt proteins in Spodoptera frugiperda. Insect Sci. 28: 627–638. https://doi.org/10.1111/1744-7917.12838

Gui FR, Lan TM, Zhao Y, Guo W, Dong Y, Fang DM, Liu H, Li HM, Wang HL, Hao RS, et al. 2022. Genomic and transcriptomic analysis unveils population evolution and development of pesticide resistance in fall armyworm Spodoptera frugiperda. Protein Cell 13: 513–531. https://doi.org/10.1007/s13238-020-00795-7

Jiang NJ, Mo BT, Guo H, Yang J, Tang R, Wang CZ. 2021. Revisiting the sex pheromone of the fall armyworm Spodoptera frugiperda, a new invasive pest in South China. Insect Sci. 0: 1–4. https://doi.org/10.1111/1744-7917.12956

Jing DP, Guo JF, Jiang YY, Zhao JZ, Sethi A, He KL, Wang ZY. 2020. Initial detections and spread of invasive Spodoptera frugiperda in China and comparisons with other noctuid larvae in corn field using molecular techniques. Insect Sci. 27: 780–790. https://doi.org/10.1111/1744-7917.12700

Johnson SJ. 1987. Migration and the life history strategy of the fall armyworm, Spodoptera frugiperda in the Western Hemisphere. Int J Trop Insect Sci. 8: 543–549.

Juárez ML, Murúa MG, García MG, Ontivero M, Vera MT, Vilardi JC, Groot AT, Castagnaro AP, Gastaminza G, Willink E. 2012. Host association of Spodoptera frugiperda (Lepidoptera: Noctuidae) corn and rice strains in Argentina, Brazil, and Paraguay. J Econ Entomol. 105: 573–582. https://doi.org/10.1603/EC11184

Kalleshwaraswamy CM, Asokan R, Mahadevaswamy HM, Sharanabasappa CM. 2019. First record of invasive fall armyworm, Spodoptera frugiperda (JE Smith) (Lepidoptera: Noctuidae) on rice (Oryza sativa) from India. J Entomol Zool Stud. 7: 332–337.

Kergoat GJ, Prowell DP, Le Ru BP, Mitchell A, Dumas P, Clamens AL, Condamine FL, Silvain JF. 2012. Disentangling dispersal, vicariance and adaptive radiation patterns: A case study using armyworms in the pest genus Spodoptera (Lepidoptera: Noctuidae). Mol Phylogenet Evol. 65: 855–870. https://doi.org/10.1016/j.ympev.2012.08.006

Kim D, Paggi JM, Park C, Bennett C, Salzberg SL. 2019. Graph-based genome alignment and genotyping with HISAT2 and HISAT-genotype. Nat Biotechnol. 37: 907–915. https://doi.org/10.1038/s41587-019-0201-4

Kumar S, Stecher G, Li M, Knyaz C, Tamura K. 2018. MEGA X: molecular evolutionary genetics analysis across computing platforms. Mol Biol Evol.35: 1547–1549.

Lee G, Bo YS, Lee J, Kim H, Song JH, Lee W. 2020. First report of the fall armyworm, Spodoptera frugiperda (Smith, 1797) (Lepidoptera: Noctuidae), a new migratory pest in Korea. Korean J Appl Entomol. 59: 73–78.

Leigh JW, Bryant D. 2015. POPART: full-feature software for haplotype network construction. Methods Ecol Evol. 6: 1110–1116.

Li H, Handsaker B, Wysoker A, Fennell T, Ruan J, Homer N, Marth G, Abecasis G, Durbin R. 2009. The sequence alignment/map format and SAMtools. Bioinformatics 25: 2078–2079. https://doi.org/10.1093/bioinformatics/btp352

Li H. 2011. A statistical framework for SNP calling, mutation discovery, association mapping and population genetical parameter estimation from sequencing data. Bioinformatics 27: 2987–2993. https://doi.org/10.1093/bioinformatics/btr509

Li, H. 2013. Aligning sequence reads, clone sequences and assembly contigs with BWA-MEM. arXiv preprint 1303.3997. https://doi.org/10.48550/arXiv.1303.3997

Librado P, Rozas J. 2009. DnaSP v5: a software for comprehensive analysis of DNA polymorphism data. Bioinformatics 25: 1451–1452. https://doi.org/10.1093/bioinformatics/btp187

Love MI, Huber W, Anders S. 2014. Moderated estimation of fold change and dispersion for RNA-seq data with DESeq2. Genome Biol. 15: 1–21. https://doi.org/10.1186/s13059-014-0550-8

Machado V, Wunder M, Baldissera VD, Oliveira JV, Fiúza LM, Nagoshi RN. 2008. Molecular characterization of host strains of Spodoptera frugiperda (Lepidoptera: Noctuidae) in Southern Brazil. Ann Entomol Soc Am. 101: 619–626. https://doi.org/10.1603/0013-8746(2008)101[619:MCOHSO]2.0.CO;2

McKenna A, Hanna M, Banks E, Sivachenko A, Cibulskis K, Kernytsky A, Garimella K, Altshuler D, Gabriel S, Daly M, et al. 2010. The genome analysis toolkit: a MapReduce framework for analyzing next-generation DNA sequencing data. Genome Res. 20: 1297–1303.

Mitchell ER, McNeil JN, Westbrook JK, Silvain JF, Lalanne-Cassou B, Chalfant RB, Pair SD, Waddill VH, Sotomayor-Rios A, Proshold FI. 1991. Seasonal periodicity of fall armyworm, (Lepidoptera: Noctuidae) in the Caribbean basin and northward to Canada. J Entomol Sci. 26: 39–50. https://doi.org/10.18474/0749-8004-26.1.39

Murúa MG, Nagoshi RN, dos Santos DA, Hay-Roe MM, Meagher RL, Vilardi JC. 2015. Demonstration using field collections that Argentina fall armyworm populations exhibit strain-specific host plant preferences. J Econ Entomol. 108: 2305–2315. https://doi.org/10.1093/jee/tov203

Nagoshi RN, Goergen G, Koffi D, Agboka K, Adjevi AKM, Plessis HD, Van den Berg J, Tepa-Yotto GT, Winsou JK, Meagher RL, et al. 2022. Genetic studies of fall armyworm indicate a new introduction into Africa and identify limits to its migratory behavior. Sci Rep. 12: 1941. https://doi.org/10.1038/s41598-022-05781-z

Nagoshi RN, Goergen G, Plessis HD, Van den Berg J, Meagher RJ. 2019. Genetic comparisons of fall armyworm populations from 11 countries spanning sub-Saharan Africa provide insights into strain composition and migratory behaviors. Sci Rep. 9: 8311. https://doi.org/10.1038/s41598-019-44744-9

Nagoshi RN, Htain NN, Boughton D, Zhang L, Xiao YT, Nagoshi BY, Mota-Sanchez D. 2020. Southeastern Asia fall armyworms are closely related to populations in Africa and India, consistent with common origin and recent migration. Sci Rep. 10: 1421. https://doi.org/10.1038/s41598-020-58249-3

Nagoshi RN, Koffi D, Agboka K, Tounou KA, Meagher RL. 2017. Comparative molecular analyses of invasive fall armyworm in Togo reveal strong similarities to populations from the eastern United States and the Greater Antilles. PLoS ONE 12: e0181982. https://doi.org/10.1371/journal.pone.0181982

Nagoshi RN, Meagher RL, Jenkins DA. 2010. Puerto Rico fall armyworm has only limited interactions with those from Brazil or Texas but could have substantial exchanges with Florida populations. J Econ Entomol. 103: 360–367. https://doi.org/10.1603/EC09253

Nagoshi RN, Meagher RL, Kathy F, Jeffrey G. RyanJ, Juan L, Armstrong JS, David BG, Chris S, Rogers LB. 2008. Using haplotypes to monitor the migration of fall armyworm (Lepidoptera: Noctuidae) corn-strain populations from Texas and Florida. J Econ Entomol.101: 742–749. https://doi.org/10.1093/jee/101.3.742

Nagoshi RN, Meagher RL, Nuessly G, Hall DG. 2006. Effects of fall armyworm (Lepidoptera: Noctuidae) interstrain mating in wild populations. Environ Entomol. 35: 561–568. https://doi.org/10.1603/0046-225X-35.2.561

Nagoshi RN, Meagher RL. 2004. Behavior and distribution of the two fall armyworm host strains in Florida. Fla Entomol. 87: 440–449. https://doi.org/10.1653/0015-4040(2004)087[0440:BADOTT]2.0.CO;2

Nagoshi RN, ROSAS-GARCIA NM, Meagher RL, Fleischer SJ, Westbrook JK, Sappington TW, Mirian HR, Thomas JMG, Murúa GM. 2015. Haplotype profile comparisons between Spodoptera frugiperda (Lepidoptera: Noctuidae) populations from Mexico with those from Puerto Rico, South America, and the United States and their implications to migratory behavior. J Econ Entomol. 108: 135–144. https://doi.org/10.1093/jee/tou044

Nagoshi RN, Silvie P, Meagher RL, Lopez J, Machado V. 2007. Identification and comparison of fall armyworm (Lepidoptera: Noctuidae) host strains in Brazil, Texas, and Florida. Ann Entomol Soc Am. 100: 394–402. https://doi.org/10.1603/0013-8746(2007)100[394:IACOFA]2.0.CO;2

Nagoshi RN, Silvie P, Meagher RL. 2007. Comparison of haplotype frequencies differentiate fall armyworm (Lepidoptera: Noctuidae) corn-strain populations from Florida and Brazil. J Econ Entomol. 100: 954–961. https://doi.org/10.1093/jee/100.3.954

Nagoshi RN. 2010. The fall armyworm triose phosphate isomerase (Tpi) gene as a marker of strain identity and interstrain mating. Ann Entomol Soc Am. 103: 283–292. https://doi.org/10.1603/AN09046

Nagoshi RN. 2012. Improvements in the identification of strains facilitate population studies of fall armyworm subgroups. Ann Entomol Soc Am. 105: 351–358. https://doi.org/10.1603/AN11138

Nagoshi RN. 2019. Evidence that a major subpopulation of fall armyworm found in the Western Hemisphere is rare or absent in Africa, which may limit the range of crops at risk of infestation. PLoS ONE 14: e0208966. https://doi.org/10.1371/journal.pone.0208966

Nam K, Nhim S, Robin S, Bretaudeau A, Nègre N, d’Alençon E. 2020. Positive selection alone is sufficient for whole genome differentiation at the early stage of speciation process in the fall armyworm. BMC Evol Biol. 20: 152. https://doi.org/10.1186/s12862-020-01715-3

Pashley DP, Hammond AM, Hardy TN. 1992. Reproductive isolating mechanisms in fall armyworm host strains (Lepidoptera: Noctuidae). Ann Entomol Soc Am. 85: 400–405. https://doi.org/10.1093/aesa/85.4.400

Pashley DP, Johnson SJ, Sparks AN. 1985. Genetic population structure of migratory moths: the fall armyworm (Lepidoptera: Noctuidae). Ann Entomol Soc Am. 78: 756–762. https://doi.org/10.1093/aesa/78.6.756

Pashley DP, Martin JA. 1987. Reproductive incompatibility between host strains of the fall armyworm (Lepidoptera: Noctuidae). Ann Entomol Soc Am. 80: 731–733. https://doi.org/10.1093/aesa/80.6.731

Pashley DP. 1986. Host-associated genetic differentiation in fall armyworm (Lepidoptera: Noctuidae): a sibling species complex?. Ann Entomol Soc Am. 79: 898–904. https://doi.org/10.1093/aesa/79.6.898

Pavlidis P, Živkovic D, Stamatakis A, Alachiotis N. 2013. SweeD: likelihood-based detection of selective sweeps in thousands of genomes. Mol Biol Evol. 30: 2224–2234. https://doi.org/10.1093/molbev/mst112

Pertea M, Pertea GM, Antonescu CM, Chang TC, Mendell JT, Salzberg SL. 2015. StringTie enables improved reconstruction of a transcriptome from RNA-seq reads. Nat Biotechnol. 33: 290–295. https://doi.org/10.1038/nbt.3122

Prowell DP, McMichael M, Silvain JF. 2004. Multilocus genetic analysis of host use, introgression, and speciation in host strains of fall armyworm (Lepidoptera: Noctuidae). Ann Entomol Soc Am. 97: 1034–1044. https://doi.org/10.1603/0013-8746(2004)097[1034:MGAOHU]2.0.CO;2

Purcell S, Neale B, Todd-Brown K, Thomas L, Ferreira MA, Bender D, Maller J, Sklar P, de Bakker PI, Daly MJ, et al. 2007. PLINK: a tool set for whole-genome association and population-based linkage analyses. Am J Hum Genet. 81: 559–575. https://doi.org/10.1086/519795

Schlum KA, Lamour K, de Bortoli CP, Banerjee R, Emrich SJ, Meagher R, Pereira E, Murua MG, Sword GA, Tessnow AE, et al. 2021. Whole genome comparisons reveal panmixia among fall armyworm (Spodoptera frugiperda) from diverse locations. BMC Genomics 22: 179. https://doi.org/10.1186/s12864-021-07492-7

Schöfl G, Heckel DG, Groot AT. 2009. Time-shifted reproductive behaviours among fall armyworm (Noctuidae: Spodoptera frugiperda) host strains: evidence for differing modes of inheritance. J Evol Biol. 22: 1447–1459. https://doi.org/10.1111/j.1420-9101.2009.01759.x

Shan YX, Jin MH, Chakrabarty S, Yang B, Li Q, Cheng Y, Zhang L, Xiao YT. 2022. Sf-FGFR and Sf-SR-C are not the receptors for Vip3Aa to exert insecticidal toxicity in Spodoptera frugiperda. Insects 13: 547. https://doi.org/10.3390/insects13060547

Song YF, Yang XM, Zhang HW, Zhang DD, Wu KM. 2021. Interference competition and predation between invasive and native herbivores in maize. J Pest Sci. 94: 1053–1063.

Sparks AN. 1979. A review of the biology of the fall armyworm. Fla Entomol. 62: 82–87. https://doi.org/10.2307/3494083

Su XN, Li CY, Xu YJ, Huang SH, Liu WL, Liao ZX, Zhang YP. 2022. Feeding preference and adaptability of fall armyworm Spodoptera frugiperda on five species of host plants and six weeds. J Environ Entomol. 44: 263–272. [in Chinese]

Sun XX, Hu CX, Jia HR, Wu QL, Shen XJ, Zhao SY, Jiang YY, Wu KM. 2021. Case study on the first immigration of fall armyworm, Spodoptera frugiperda invading into China. J Integr Agr. 20: 664–672. https://doi.org/10.1016/S2095-3119(19)62839-X

Swamy HMM, Asokan R, Kalleshwaraswamy CM, Deshmukh SS, Prasad YG, Maruthi MS, Shashank PR, Devi NI, Surakasula A, Adarsha S, et al. 2018. Prevalence of “R” strain and molecular diversity of fall army worm Spodoptera frugiperda (J.E.Smith) (Lepidoptera: Noctuidae) in India. Indian J Entomol. 80: 544–553. 10.5958/0974-8172.2018.00239.0

Tay WT, Rane RV, Padovan A, Walsh TK, Elfekih S, Downes S, Nam K, d’Alençon E, Zhang JP, Wu YD, et al. 2022. Global population genomic signature of Spodoptera frugiperda (fall armyworm) supports complex introduction events across the Old World. Commun Biol. 5: 297. https://doi.org/10.1038/s42003-022-03230-1

Withers AJ, de Boer J, Chipabika G, Zhang L, Smith JA, Jones CM, Wilson K. 2021. Microsatellites reveal that genetic mixing commonly occurs between invasive fall armyworm populations in Africa. Sci Rep. 11: 20757. https://doi.org/10.1038/s41598-021-00298-3

Withers AJ, Rice A, de Boer J, Donkersley P, Pearson AJ, Chipabika G, Karangwa P, Uzayisenga B, Mensah BA, Mensah SA, et al. 2022. The distribution of covert microbial natural enemies of a globally invasive crop pest, fall armyworm, in Africa: enemy-release and spillover events. J Anim Ecol. 00: 1–16. https://doi.org/10.1111/1365-2656.13760

Wu KM. 2020. Management strategies of fall armyworm (Spodoptera frugiperda) in China. Plant Prot. 46: 1–5. [in Chinese]

Xiao HM, Ye XH, Xu HX, Mei Y, Yang Y, Chen X, Yang YJ, Liu T, Yu YY, Yang WF, et al. 2020. The genetic adaptations of fall armyworm Spodoptera frugiperda facilitated its rapid global dispersal and invasion. Mol Ecol Resour. 20: 1050–1068. https://doi.org/10.1111/1755-0998.13182

Yang F, Morsello S, Head GP, Sansone C, Huang FN, Gilreath RT, Kerns DL. 2018. F2 screen, inheritance and cross-resistance of field-derived Vip3A resistance in Spodoptera frugiperda (Lepidoptera: Noctuidae) collected from Louisiana, USA. Pest Manag Sci. 74: 1769–1778. https://doi.org/10.1002/ps.4805

Zhang L, Liu B, Zheng WG, Liu CH, Zhang DD, Zhao SY, Li ZY, Xu PJ, Wilson K, Withers A, et al. 2020. Genetic structure and insecticide resistance characteristics of fall armyworm populations invading China. Mol Ecol Resour. 20: 1682–1696. https://doi.org/10.1111/1755-0998.13219

